# Trastuzumab Deruxtecan Combination Strongly Enhances Responses and Overcomes Sotorasib Resistance in *KRAS*^G12C^-Mutant NSCLC

**DOI:** 10.1101/2024.12.17.628751

**Authors:** Hilal Ozakinci, Bina Desai, Denise Kalos, Dung-Tsa Chen, Menkara Henry, Hitendra Solanki, Theresa A. Boyle, Humberto E. Trejo Bittar, Gina S. Nazario, Eric B. Haura, Andriy Marusyk, Bruna Pellini

## Abstract

**Introduction:** Recent advances in the treatment of *KRAS*-mutant non-small cell lung cancer (NSCLC) have led to the development of KRAS^G12C^ inhibitors, such as sotorasib and adagrasib. However, resistance and disease progression remain significant challenges. In this study, we investigated the therapeutic potential of combining trastuzumab deruxtecan (T-DXd), an anti-HER2 antibody-drug conjugate, with sotorasib in *KRAS^G12C^*-mutant NSCLC, while also evaluating HER2 expression in NSCLC samples.

**Methods:** The HER2 expression dependence of xenograft responses to sotorasib, T-DXd, and their combination was evaluated in therapy-naïve and sotorasib-treated tumors by immunohistochemistry (IHC). Also, we analyzed 191 clinical (pre- or on-treatment) and rapid autopsy (post-treatment) samples from 31 patients with driver-positive and driver-negative advanced stage NSCLC, assessing HER2 expression using interpretation guidelines developed for breast cancer (BC) and gastroesophageal adenocarcinoma (GEA).

**Results:** In the majority of preclinical models, including sotorasib-resistant tumors, the sotorasib-T-DXd combination induced stronger and more durable responses compared to monotherapies. The strong effect of the combination therapy was likely attributable to sotorasib-induced adaptive HER2 upregulation; stronger HER2 expression in sotorasib-treated tumors was linked with stronger responses. Although HER2 expression was higher in samples from patients with *KRAS^G12C^*-mutant NSCLC compared to NSCLC with other driver mutations or no drivers, the difference was not statistically significant. HER2 IHC score discrepancy was also observed between BC and GEA interpretation guidelines.

**Conclusions:** Our results support the potential clinical utility of the sotorasib-T-DXd combination, including tumors with intrinsic and acquired resistance to sotorasib monotherapy. Since the strengths of the responses depend on HER2 expression levels, successful clinical implementation necessitates optimizing patient selection. Our results highlight the complexities of accurate HER2 interpretation in NSCLC and highlight the need for standardized testing methods.

## 1. Introduction

Precision treatment of cancers harboring oncogenic *KRAS* mutation has been revitalized with enthusiasm following the discovery of new efficacious small molecules that target mutant *KRAS*^G12C^. The lead clinical compounds sotorasib (AMG-510) and adagrasib (MRTX849) demonstrated favorable effects with approximately 40% response rates, however most patients develop disease progression within 5 to 6 months.^1–3^ Given the limited duration of disease control seen with KRAS^G12C^-inhibitors (KRAS^G12C^i) monotherapy, these agents are only approved as second-line treatment, and there is a strong unmet need to develop new treatment strategies for patients with *KRAS*^G12C-^mutant non-small cell lung cancer (NSCLC). Due to the distinct features of RAS signaling, this need is unlikely to be solved through advances in KRAS^G12C^i design. Therefore, improving the response to KRAS^G12C^i calls for the development of combination therapy strategies. The main approach in the combination therapies space has been the search for the molecular mechanisms responsible for KRAS^G12C^i resistance and combining KRAS^G12C^i with pharmacological inhibitors of these mechanisms of resistance, or with classic chemotherapeutic agents or immune checkpoint inhibitors.^4–6^ Extensive research in this area led to the identification of a large and growing list of genomic and non-genomic mechanisms of resistance, including many potentially targetable ones.^7–9^

Therapeutic success of resistance mechanism-focused combination therapies faces several major challenges. These challenges include intra-tumor heterogeneity (the presence of subpopulations with different resistance mechanisms within the same tumor)^10^, complexity of resistance phenotypes (contribution of multiple mechanisms to a given resistance phenotype), and resistance-promoting interactions with the tumor microenvironments where resistance phenotypes are shaped by combination of both cell-intrinsic and cell-extrinsic mechanisms.^8,9,11^ Therefore, combination therapies focused on individual resistance mechanisms are unlikely to provide a substantial improvement in responses to RAS targeting, colling for consideration of orthogonal approaches.

Antibody-drug conjugates (ADC), a class of drugs that combines an antibody directed against a specific epitope with a non-specific cytotoxic payload, offer the potential solution to the challenges faced by approaches focused on specific resistance mechanisms.^12,13^ The effect of antibody-based therapies is independent of whether the epitope is essential for KRAS^G12C^i resistance, sidestepping the challenge of the complexity of resistance phenotypes. As long as the targeted epitope is expressed by a substantial proportion of tumor cells, the use of an ADC with a payload capable of inducing bystander damage might alleviate the challenge of intra-tumor heterogeneity. Finally, ADCs potentially enable achieving high local concentrations of effective cytotoxic therapies while avoiding their systemic side effects.

Fam-Trastuzumab Deruxtecan (T-DXd), an ADC that combines trastuzumab, an antibody that targets HER2 protein (encoded by the *ERBB2* gene), with the topoisomerase 1 inhibitor deruxtecan, connected by through a tetrapeptide-based cleavable linker, offers a particularly attractive option. In vitro and in vivo studies have observed that the activation of ERBB signaling aids *KRAS*^G12C^ signaling leading to tumor progression.^14–16^ Importantly, in contrast to trastuzumab, T-DXd responses are not limited to tumors with high HER2 expression, as strong and lasting responses have also been documented in breast cancers with low HER2 expression^17^. Importantly, similar to the effects observed in EGFR^18^ and ALK^19^ targeted therapies, KRAS inhibition can lead to a substantial increase in HER2 expression, reflective of both adaptive induction and selective advantage of HER2 overexpressing cells.^14^ Therefore, KRAS^G12C^i treatment can be expected to potentiate responses to T-DXd. Finally, a combination of T-DXd with the inhibition of primary signaling driver has induced remarkably strong responses in animal models of *ALK*-mutant NSCLC.^20^

Based on these premises, we investigated the utility of T-DXd - sotorasib combination in a panel of mouse models of *KRAS*^G12C^-mutant NSCLC and evaluated HER2 expression. To explore the translational potential of findings from mouse models, we examined HER2 expression in clinical (pre-treatment and on-treatment) and rapid autopsy (post-treatment) tumor samples from patients with NSCLC. Given the lack of a specific interpretation guidelines for HER2 expression in NSCLC, HER2 expression in NSCLC was tested with two existing interpretation guidelines developed for therapeutic decision-making in clinical practice for breast cancer (BC) and gastric/gastroesophageal adenocarcinoma (GEA) and compared their results in clinical and rapid autopsy samples from primary and metastatic tumors.

## 2. Materials and Methods

### 2.1. Xenograft studies

For the initiation of xenograft tumors, one million cells were suspended in 100 µl of 1:1 RPMI and BME-Type 3 (R&D Systems, # 36-320-1002P) per implantation. Cells were implanted via two contralateral subcutaneous injections in NOD-Scid IL2Rgnull (NSG) recipient mice aged between 4 to 6 weeks. After the tumors reached 3-4 mm in diameter, mice were randomly assigned to the control or treatment groups. Baseline tumor measurements were recorded before initiation of treatment. 50 mg/Kg of sotorasib (suspended in 0.25% of methylcellulose and 0.05% Tween 80 in water) was administered via daily oral gavages, and 10 mg/Kg of T-DXd was administered once every two weeks via intraperitoneal injections. Tumor diameters were recorded weekly using an electronic caliper; tumor volumes were calculated with the assumption of the tumor’s spherical shape. All animal experiments were approved and performed by the Institutional Animal Care and Use Committee (IACUC) guidelines of the H. Lee Moffitt Cancer Center. Animals were produced in the in-house breeding colony, with breeders purchased from Jackson Laboratory and housed in a specific pathogen-free vivarium accredited by the Association for the Assessment and Accreditation of Laboratory Animal Care (AAALAC), which includes temperature, humidity control, and a 12-hour light/12-hour dark cycle.

### 2.2. Study Population

The Rapid Tissue Donation (RTD) program at Moffitt Cancer Center has enabled patients with advanced stage lung cancer to donate tumor tissue samples, as well as body fluids, for research after death.^21^ As of June 2024, 99 patients were consented for the RTD and autopsies were performed in 58 patients with a mean of 14 hours between death and collection of paired tissue samples processed as formalin-fixed paraffin-embedded (FFPE) and frozen tissue. During the patient/sample selection process for this study, the following inclusion criteria were applied: i) patients diagnosed with adenocarcinoma of the lung or NSCLC, NOS and ii) samples showing an adequate quantity of viable tumor cells (tumor cell percentage ≥10% and tumor cell count ≥100). Additionally, patients or samples exhibiting the following features were excluded from the study: i) NSCLC with small cell lung cancer transformation, and ii) samples containing only carcinomatosis. If multiple FFPE samples from same tumor were available, only one FFPE with the highest tumor volume was chosen for downstream analysis. In addition to the autopsy samples, all available clinical samples, which were collected during patient care for diagnostic purposes, were evaluated for adequacy. Clinical samples meeting the criteria of a tumor cell percentage ≥10% and a tumor cell count ≥100 were included in the study. A total of 161 autopsy samples and 30 clinical samples from 31 patients with NSCLC were analyzed. All autopsy samples were post-treatment, whereas 16 of the 30 clinical samples were pre-treatment, and the remaining 14 were on-treatment. Both pre-treatment clinical and post-treatment autopsy samples from same tumor site were available for 9 patients. Clinical characteristics were extracted from electronic medical records.

### 2.3. HER2 Immunohistochemistry

In xenograft tumors, immunohistochemistry (IHC) was performed on 5 μm FFPE tumor tissues. Antigen retrieval was performed in citrate buffer (pH 6.2), followed by incubation with 3% hydrogen peroxide in methanol. Slides were blocked by 10% goat serum, then stained with primary anti-HER2 antibody (cell signaling #4290 - D8F12, 1:400) for 1 hour and secondary anti-rabbit biotinylated antibody (Vector Labs, BA-1000, 1:100) for 45 mins at room temperature. Vectastain ABC peroxidase reagent (Vector Laboratories, PK-6100) with DAB as a colorimetric substrate was used for the antibody detection followed by hematoxylin counterstain. In human samples, IHC was performed using FDA-approved in vitro diagnostic (IVD) clone 4B5 on the Ventana BenchMark ULTRA automated system (Catalog number: 790-2991, Roche Diagnostics, Indianapolis, IN) as per manufacturer’s protocol.^22^ Additionally, the research use only (RUO) clone EP1045Y (ab134182, dilution 1:200, incubation 15 minutes, Abcam, Cambridge, UK) was used to analyze 34 human tumor samples on the Leica Bond RX automated system (Leica Biosystems, Buffalo Grove, IL) as per manufacturer’s protocol^23^. IHC results were interpretated according to American Society of Clinical Oncology (ASCO) / College of American Pathologists (CAP) HER2 interpretation guidelines for BC (March 2023), and GEA (June 2017) (**Supplementary Table 1**). ^24,25^

### 2.4. HER2 Dual In-Situ Hybridization

The Ventana HER2/neu Dual ISH DNA Probe Cocktail assay was performed on samples with discordant IHC results or samples with an IHC score of 2+ with both (BC and GEA) interpretation guidelines. Manufacture’s validated and recommended protocol (Catalog number:760-6072, Roche Diagnostics, Indianapolis, IN) was used.^26^ At least 20 tumor cells per slide were manually counted by the pathologist. ISH results were documented in accordance with ASCO/CAP guidelines for BC and GEA.^25,27^

### 2.5. Statistical Analysis

Descriptive statistics were used to summarize sample characteristics (e.g., tissue type, HER2 IHC score). Agreement percentages and McNemar’s Chi-squared test were used to compare HER2 scoring interpretation guidelines in sample level and patient level. Differences in expression based on driver or type of tissue were tested using Fisher’s exact or a two-sample test of equality of proportions. All p-values are unadjusted.

## 3. Results

### 3.1. T-DXd enhances sotorasib sensitivity and overcomes sotorasib resistance in the H358 xenograft model of KRAS^G12C^-mutant NSCLC

To assess the utility of T-DXd in potentiating responses to sotorasib in vivo, we started with a common experimental model of *KRAS*^G12C^-mutant NSCLC H358 cell line. Upon subcutaneous implantation into immune-deficient NSG mice, H358 cells form rapidly growing tumors that display abundant HER2 expression primarily localized to the plasma membrane (**Figure 1A**, **left**). Sotorasib treatment of H358 xenograft tumors induced a substantial remission, with an approximately 10-fold reduction in tumor volume. However, after 10-12 weeks of continuous treatment, the tumors have relapsed (**Figure 1B**, **Figure S1A**), recapitulating a pattern seen in a significant subset of human patients with sotorasib-responsive *KRAS*^G12C^-mutant tumors. Assessment of tumors that have progressed on sotorasib monotherapy revealed that, consistent with our prior in vitro observations^14^, sotorasib treatment further enhanced HER2 expression levels (**Figure 1A**, **right**), bolstering the rationale to explore the sotorasib-T-DXd combination. Whereas T-DXd monotherapy failed to induce regression of H358 tumors, it potently enhanced responses to sotorasib, driving an approximately 1000-fold reduction in tumor volume and preventing the relapse at least up to 16 weeks on-treatment (**Figure 1B**, **Figure S1A**).

**Figure 1:**
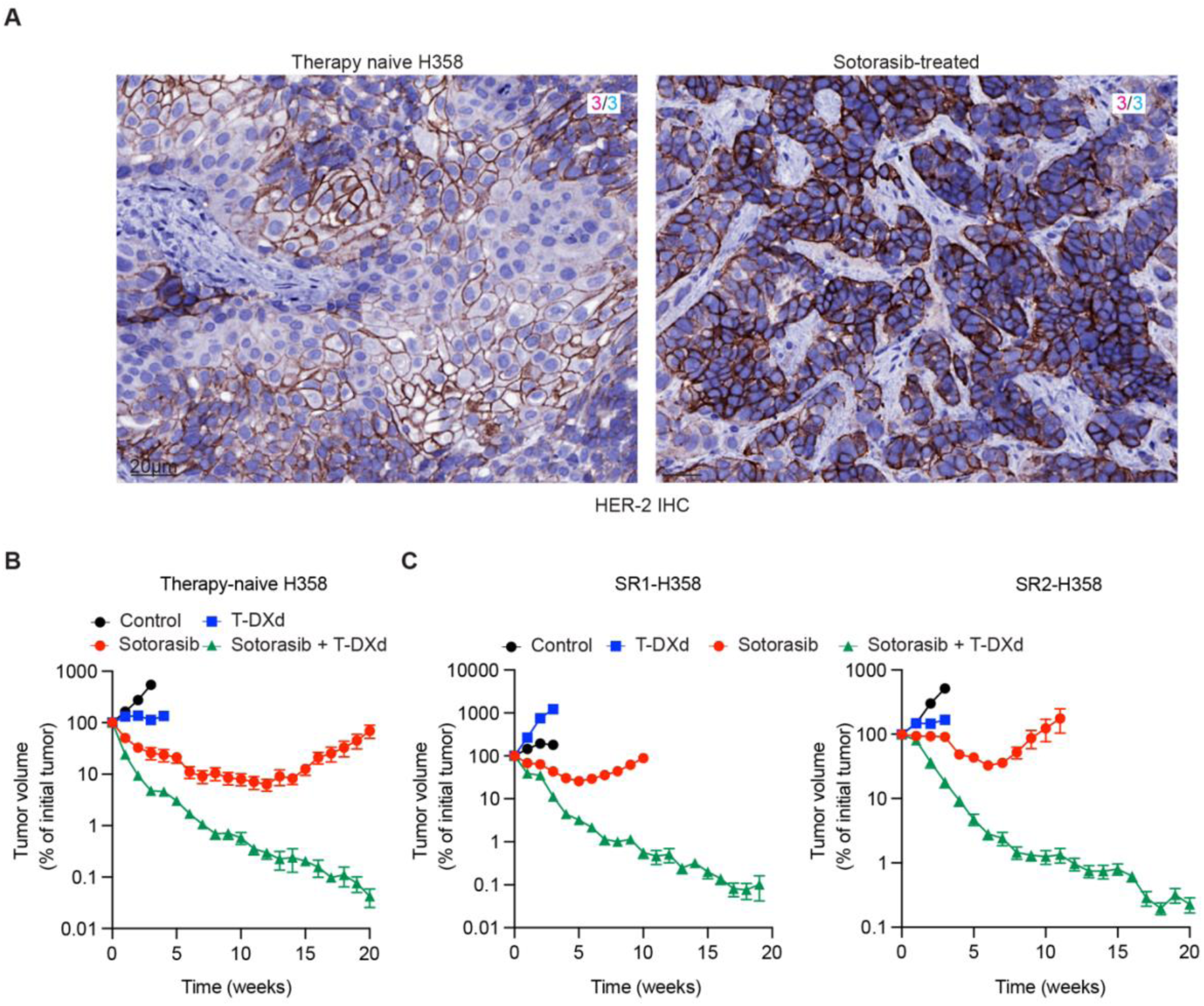
TDX-d enhances sotorasib sensitivity of therapy-naïve and sotorasib-resistant (SR) H358 xenograft tumors. **A**. Representative images of HER2 IHC staining of therapy-naïve and sotorasib-treated tumors. Both tumors are showing high HER2 expression with a IHC score of 3+. Scale bars, 20μm. **B**. Volumetric response dynamics of H358 xenograft tumors to vehicle (control, N=6), 50 mg/kg sotorasib (N=10), 10 mg/kg T-DXd (N=8), and sotorasib/T-DXd combinations (N=10). Error bars represent SEM. **C**. Volumetric response dynamics of secondary transplants of two independent H358 tumors that have previously progressed on sotorasib to vehicle (control, N=8), 50 mg/kg sotorasib (N=10), 10 mg/kg T-DXd (N=6 for SR1, N=8 for SR2), and sotorasib/T-DXd combinations (N=10). Error bars represent SEM.

Given the remarkable enhancement of sotorasib sensitivity, we sought to examine the effect of T-DXd treatment on tumors that have developed sotorasib resistance (SR). To this end, we have isolated cells from two independent H358 xenograft tumors that have relapsed on sotorasib monotherapy (**SR1 and SR2**, **Figure S1B**), expanded these cells ex-vivo, and initiated cohorts of secondary transplants. Sotorasib monotherapy induced regression in both models, indicating a partial re-sensitization in the absence of therapy; however, the responses were more shallow and shorter-lasting compared to the responses of therapy naïve H358 xenografts (**Figure 1C, Figure S1C**). In contrast, even though T-DXd monotherapy failed to induce regression in either of the SR models, the sotorasib-T-DXd combination induced a strong and durable regression compatible with the regression observed in therapy-naïve xenografts (**Figure 1C, Figure S1C**). Therefore, T-DXd might be capable of overcoming, sotorasib resistance, restoring sensitivity to KRAS inhibition.

### 3.2. Sensitivity of KRAS^G12C^-mutant NSCLC xenografts to sotorasib-T-DXd combination correlates with HER2 expression

Given the substantial inter-tumor variability of *KRAS*^G12C^-mutant NSCLC, we assess the generalizability of the ability of T-DXd to enhance sotorasib sensitivity. To this end, we examined therapeutic responses in a panel of 8 additional xenograft models of the disease. While none of these additional models displayed the strong sotorasib sensitivity observed in H358 xenografts, 5/8 displayed either a stabilization of the tumor volume or a weak response, followed by progression within 1-5 weeks (**Figure S2**). In 6/8 models, T-DXd monotherapy inhibited tumor growth but failed to induce regression. In contrast, consistent with sotorasib/T-DXd combination sensitivity in the H358 xenografts, sotorasib-T-DXd combination induced remissions in 7/8 of the additional *KRAS*^G12C^ xenografts (**Figure S2**).

Given that T-DXd sensitivity requires HER2 expression, we sought to examine the relationship between HER2 expression levels and the sensitivity to the sotorasib/T-DXd combination. IHC is the standard method to assess HER2 expression in clinical diagnostics. However, clinical guidelines for IHC-based assessment of HER2 expression in NSCLC have not yet been developed.^28^ Therefore, we elected to assess the HER2 expression using two IHC interpretation guidelines developed for therapeutic decision-making in clinical practice for GEA^29^ and BC^27^.

Even though sotorasib treatment has enhanced the percentage of tumor cells showing membranous HER2 expression in H358 tumors (both therapy-naïve and sotorasib-treated tissues were processed and stained in the same batch), the IHC scores with both interpretation guidelines (GEA and BC) fell into same group as 3+, indicating a limited resolution of interpretation guidelines to detect expression differences. Analyses of therapy-naïve and post-sotorasib tissues collected upon relapse revealed substantial variability in both baseline and sotorasib-induced HER2 expression, with scores ranging from 0 to 3+. At least in some models, sotorasib treatment resulted in enhanced HER2 expression that could be captured by IHC interpretation guidelines (**Figure 2A**). While for therapy-naïve tumors, GEA and BC interpretation guidelines yielded identical scores, some divergence was observed for sotorasib-treated tumors (**Figure 2A**, **asterisks**).

**Figure 2:**
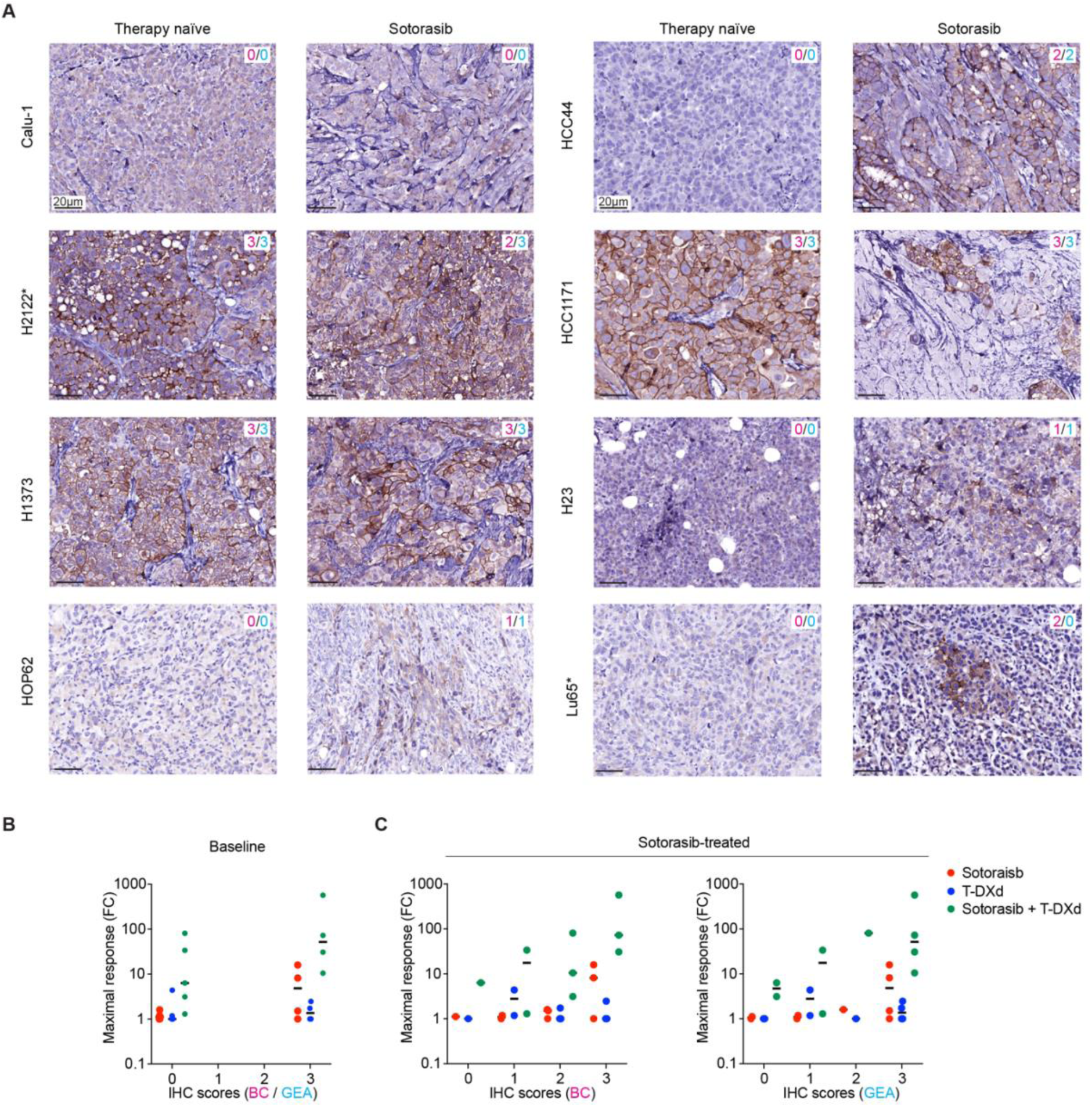
**A.** Representative images of HER2 IHC of therapy-naïve and sotorasib-treated tumors from the indicated xenograft models of *KRAS*^G12C^-mutant NSCLC. HER2 IHC scoring of whole tumor tissue based on the interpretation guidelines for BC (pink)/GEA (blue) are indicated on the top right. **B, C.** Maximal magnitude of tumor regression in response to the indicated therapies as a function of the IHC-based scoring (BC and GEA) of therapy-naïve (**B**) and sotorasib-treated (**C**) tumors from the nine *KRAS*^G12C^-mutant xenograft models. IHC scores using both the BC and GEA interpretation guidelines were concordant in therapy-naïve samples, but discrepancies between the two interpretation guidelines were observed in sotorasib-treated tumors.

To assess the relationship between HER2 expression and therapeutic sensitivities, we plotted the maximal magnitude of tumor regression (expressed in fold change) as a function of IHC interpretation guidelines. No apparent relation was observed for tumor sensitivities to sotorasib and T-DXd monotherapies (**Figure 2B**). Tumors with high HER2 baseline expression (score of 3+) displayed stronger responses to the combination therapy, although the difference was not statistically significant. Plots of the relationship between HER2 expression in sotorasib-treated tumors and therapeutic responses to the sotorasib-T-DXd combination suggested a stronger correlation, although the limited number of models and substantial variability in tumor responses precluded inferences of statistical significance (**Figure 2C**). In summary, these analyses support the expected dependence of sotorasib-T-DXd responses on HER2 expression levels while highlighting the limitations of the available HER2 IHC interpretation guidelines for BC and GEA.

### 3.3. Patient characteristics

Given the observed enhancement in T-DXd sensitivity in a panel of xenograft models of *KRAS*^G12C^-mutant NSCLC and the relationship between the combination sensitivity and HER2 expression, we evaluated HER2 expression in a cohort of thirty-one NSCLC patients. The study patients had a median age of 62 years (range 47 to 86 years). Most patients were female (58%) and White (97%). All patients had stage IV NSCLC, with adenocarcinoma being the predominant histology (97%). High PD-L1 expression (≥50%) was observed in 26% of the tested samples. Driver genomic alterations were detected in 87% of patients during clinical care, with *KRAS* mutations being the most common (55%), followed by *ALK* (13%), *EGFR* (10%), *MET* (3%), *NRAS* (3%) and *ROS1* (3%). *HER2* mutation or amplification was not detected in the tested samples. At baseline, 71% of patients had an ECOG performance status of 1, and 61% of the patients received targeted therapy. The most common tumor sites in the 161 autopsy samples were the lungs (29%), lymph nodes (25%), and liver (13%), with 6% of tumors collected from the brain (**Table 1**).

**Table 1.** Demographics and clinical features of patients.

| Characteristics | NSCLC (n=31) |
| --- | --- |
| Age, years, median (range) | 62 (47-86) |
| Female, No. (%) | 18 (58) |
| Race, No. (%) |  |
| White | 30 (97) |
| Black | 1 (3) |
| Disease stage at enrollment, No. (%) |  |
| IV | 31 (100) |
| Diagnosis |  |
| Adenocarcinoma | 30 (97) |
| NSCLC, NOS | 1 (3) |
| PD-L1 protein expression, No. (%) |  |
| <1% | 12 (39) |
| ≥1 to 49% | 2 (6) |
| ≥50% | 8 (26) |
| Not available | 9 (29) |
| Driver genomic alteration |  |
| KRAS | 17 (55) |
| ALK | 4 (13) |
| EGFR | 3 (10) |
| ROS1 | 1 (3) |
| MET | 1 (3) |
| NRAS | 1 (3) |
| No driver | 4 (13) |
| ECOG PS at baseline, No. (%) |  |
| 0 | 7 (23) |
| 1 | 22 (71) |
| 2 | 2 (6) |
| Type of systemic therapy, No. (%) |  |
| Platinum-based chemotherapy | 29 (94) |
| Immunotherapy | 26 (84) |
| Targeted small molecules | 19 (61) |
| Tumor sites at the autopsy*, No. (%) |  |
| Lung | 47 (29) |
| Lymph nodes | 41 (25) |
| Liver | 21 (13) |
| Brain | 9 (6) |
| Others | 43 (27) |
| Tumor sites at the clinical samples, No. (%) |  |
| Lung | 11 (37) |
| Lymph nodes | 4 (13) |
| Liver | 1 (3) |
| Others | 14 (47) |

\* Autopsy samples with tumor cell percentage ≥10% and tumor cell count ≥100 are listed.

### 3.4. HER2 expression in NSCLC using ASCO/CAP interpretation guidelines developed for BC and GEA

The distribution of tumors with IHC scores of 0, 1+, 2+, and 3+ was as follows: 70%, 10%, 12%, 7.9% and 68%, 2%, 20%, 10% according to BC and GEA interpretation guidelines, respectively (**Table 2**). Among the 191 samples analyzed, increased HER2 expression (2+ or 3+) with either one of interpretation guidelines was detected in 58 (30%) samples. HER2 IHC scores were concordant for 166 (87%) samples using both interpretation guidelines (BC and GEA), the remaining 25 samples (13%) showed discordance between the two interpretation guidelines (**Table 2**, **Figure 3**). When IHC scores were grouped into three categories (3+, 2 vs 0 or 1+), 65% of patients demonstrated 100% agreement between the two interpretation guidelines across all samples. This agreement increased to 74% when the scores were categorized into two groups (2+ or 3+ vs 0 or 1+). Interestingly, nuclear HER2 expression was observed in 25 samples; in 17 of these, nuclear expression was present in tumor cells, while in the remaining 8, it was detected in adjacent non-tumoral adrenal or liver cells. Cytoplasmic staining was also noted in 17 tumoral samples with nuclear HER2 expression (**Supplementary Figure 3**). Membranous HER2 expression accompanied nuclear expression in 9 out of 17 samples. Among these, two metastatic samples from two patients with *KRAS*-mutant NSCLC had a HER2 IHC score of 2+ according to both interpretation guidelines.

**Figure 3.**
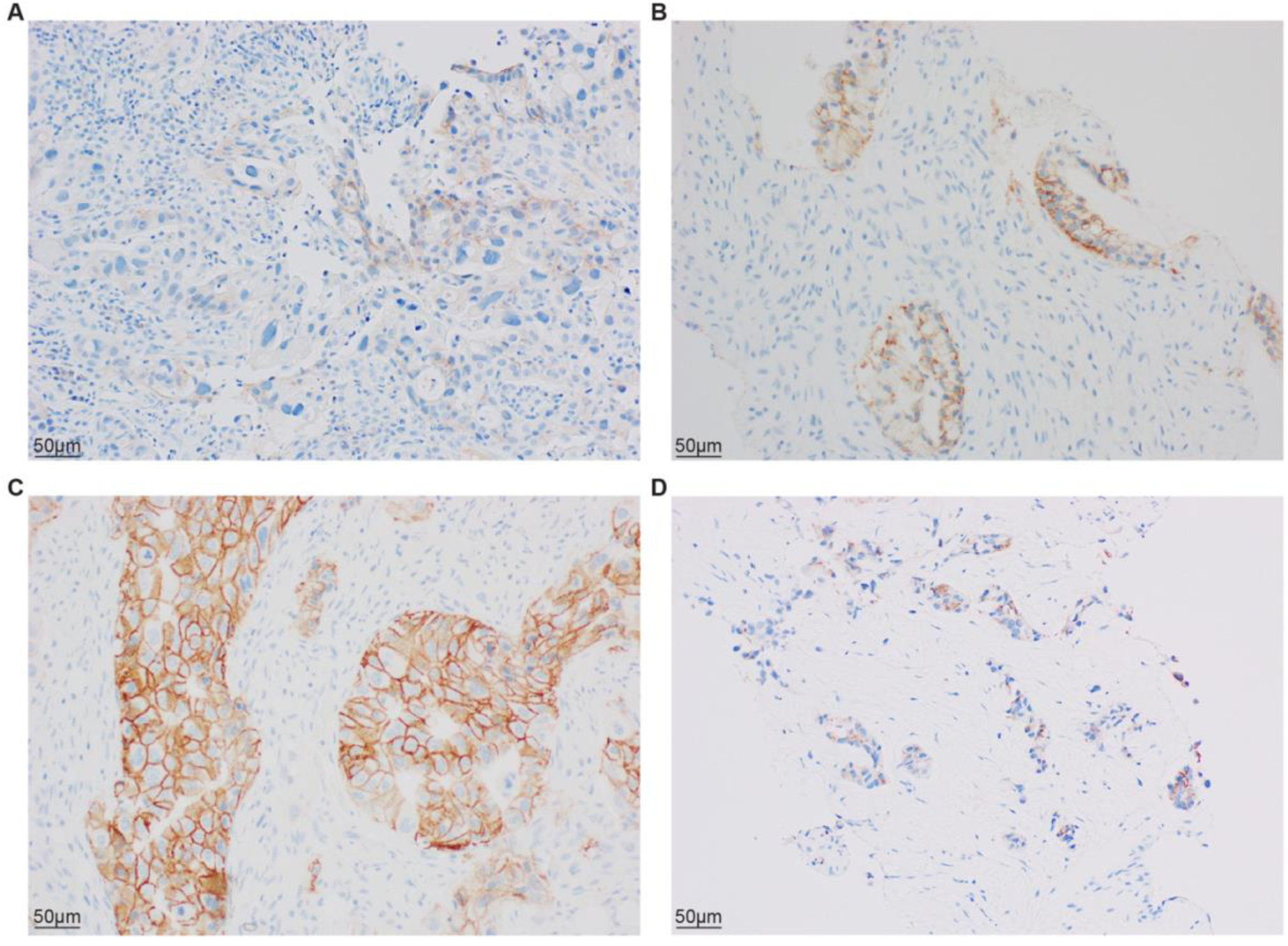
HER2 IHC interpretation according to the American Society of Clinical Oncology and College of American Pathologists (ASCO/CAP) interpretation guidelines for breast cancer (BC) vs. gastric/gastroesophageal adenocarcinoma (GEA). **A.** Section showing a HER2 IHC score of 1+, suggesting negative IHC results (x20); **B.** Section showing a HER2 IHC score of 2+, indicating equivocal IHC results (x20); **C.** Section showing a HER2 IHC score of 3+, representing a positive IHC result with high HER2 protein expression (x20). **D.** A hematoxylin and eosin (H&E) stained section shows a poorly differentiated adenocarcinoma in the adrenal gland originating from lung (x10); IHC-stained section revealing a 2+ IHC score based on the ASCO/CAP guideline for GEA, but score is 0 when using the guideline for BC(clone: 4B5,x20). This indicates that the scoring criteria can differ significantly depending on the used guideline.

**Table 2.** Comparison of HER2 IHC scores according to interpretation guidelines for BC versus GEA across all samples.

|  |  | HER2 IHC score (GEA) |  |  |  |
| --- | --- | --- | --- | --- | --- |
|  |  | 3+ | 2+ | 0 or 1+ | Total |
| HER2 IHC score (BC) | 3+ | 15 (7.9%) | 0 (0%) | 0 (0%) | 15 (7.9%) |
|  | 2+ | 5 (2.6%) | 18 (9.4%) | 0 (0%) | 23 (12%) |
|  | 0 or 1+ | 0 (0%) | 20 (10%) | 133 (70%) | 153 (80%) |
| Total |  | 20 (10%) | 38 (20%) | 133 (70%) | 191 (100%) |

### 3.5. HER2 expression in NSCLC with driver genomic alteration vs. no driver & KRAS mutation vs. other driver genomic alteration

Among the 27 patients with tumors harboring a driver mutation, 12 (44%) had at least one sample with a HER2 IHC score of 2+ or 3+ using the BC interpretation guideline, while 15 (56%) had samples with HER2 IHC score of 2+ or 3+ using the GEA interpretation guideline. In the remaining 4 patients with tumors lacking a driver mutation, 1 (25%) patient had at least one sample with a HER2 IHC score of 2+ or 3+ using the BC interpretation guideline and 2 (50%) patients had at least one sample with a HER2 IHC score of 2+ or 3+ using the GEA interpretation guideline (**Supplementary Table 2**). In 17 patients with *KRAS*-mutant tumors, 10 (59%) had at least one sample with HER2 IHC scores of 2+ or 3+ using the BC interpretation guideline, and 11 (65%) using the GEA interpretation guideline. In comparison, among 10 patients with tumors with other driver genomic alterations (*ALK, EGFR, MET, NRAS,* and *ROS1*), 2 (20%) and 4 (40%) had at least one sample with HER2 IHC scores of 2+ or 3+ using the BC and GEA interpretation guidelines, respectively (**Table 3, Supplementary Table 3**). Despite the higher HER2 expression observed in patients with *KRAS*-mutant NSCLC, there was no statistically significant difference between *KRAS*-mutant NSCLC vs. NSCLC with no driver genomic alteration (**Supplementary Table 4**) and *KRAS^G12C^*-mutant NSCLC vs. *KRAS^non-G12C^*-mutant NSCLC (**Supplementary Table 5**).

**Table 3.** Comparison of HER2 IHC scores according to GEA interpretation guideline in patients with *KRAS-*mutant *NSCLC* versus NSCLC with other drivers.

| Driver mutation | *KRAS*<br>N=17 | Other drivers*<br>N=10 | p-value |
| --- | --- | --- | --- |
| Patients with at least one sample showing 2+ and/or 3+ HER2 IHC score, No. (%) | 11 (65%) | 4 (40%) | >0.9 |

\**ALK, EGFR, MET, NRAS, ROS1*

### 3.6. HER2 expression in clinical vs. autopsy samples & pre-treatment clinical vs. post-treatment autopsy samples

HER2 IHC scores of 2+ or 3+ were observed in 28 out of 161 (17%) autopsy samples and in 10 out of 30 (33%) clinical samples using the BC interpretation guideline. With GEA interpretation guideline, these scores were found in 45 out of 161 (28%) autopsy samples and 13 out of 30 (43%) clinical samples. A significantly higher prevalence of HER2 positivity was observed in clinical samples compared to autopsy samples (p = 0.039) with BC interpretation guideline, while the difference was not statistically significant using the GEA interpretation guideline (p = 0.071). Among the 31 patients with NSCLC, both pre-treatment clinical and post-treatment autopsy samples from same tumor were available for 9 patients. When HER2 IHC scores were compared between pre-treatment clinical and post-treatment autopsy sample from same tumor site of 9 patients, the HER2 IHC score remained unchanged in four patients, decreased in three, and increased in two (**Supplementary Table 6**).

### 3.7. IVD vs RUO HER2 clones

HER2 IHC scores were assessed in 26 samples using two HER2 clones (IVD and RUO). According to the BC interpretation guideline, HER2 IHC scores of 2+ or 3+ were observed in 7.7% of samples with the IVD clone and 58% with the RUO clone. Using the GEA interpretation guideline, the IVD clone identified 12% of samples as 2+ or 3+ while the RUO clone identified 69%. These findings indicate a significantly higher HER2 IHC score with the RUO clone compared to the IVD clone, with a statistically significant difference (p<0.001) (**Supplementary Figure 4, Supplementary Table 7**).

### 3.8. *HER2* gene amplification by DISH

DISH was performed on 38 samples, which included 25 with discordant IHC results and 13 with equivocal (2+) IHC results between the BC and GEA interpretation guidelines. Among the 25 tumors with discordant IHC results, only 1 sample (score of 3+ with the GEA interpretation guideline and 2+ with the BC interpretation guideline) showed *HER2* gene amplification according to both BC and GEA DISH interpretation guidelines. The remaining 24 samples were either negative or required additional work-up, such as re-evaluation by a second pathologist or analysis of additional tumor samples, according to at least one of the BC or GEA DISH interpretation guidelines. None of the 13 samples with equivocal IHC results showed gene amplification; their DISH results were either negative or required additional work-up (**Supplementary Table 8** and **Supplementary Figure 5**).

## 4. Discussion

Our study demonstrated that HER2-targeting antibody-drug conjugate T-DXd can strongly enhance sotorasib sensitivity. This enhancement is likely attributable to the adaptive induction of HER2 expression following KRAS inhibition in *KRAS^G12C^*-mutant tumors, observed in both previous in vitro studies^14,30^ as well as in a subset of in vivo xenograft models. Importantly, the sotorasib-T-DXd combination could induce strong and durable remission in tumors with acquired sotorasib resistance. These preclinical findings have laid the groundwork for the development of a first-in-human Phase I/II clinical trial, recently approved by the Cancer Therapy Evaluation Program (CTEP) at the National Cancer Institute (NCI) titled “A Phase I/II Study Evaluating the Safety, Tolerability, and Efficacy of Sotorasib Plus Trastuzumab Deruxtecan in Patients with Advanced Non-Small Cell Lung Cancer with a KRASG12C Mutation”. This trial, currently under protocol development, will investigate the safety and efficacy of combining T-DXd with sotorasib in patients with *KRAS*^G12C^-mutant NSCLC. This represents a critical step forward in translating our preclinical findings into potential new treatment options for patients with this challenging form of lung cancer.

Given the expected dependence of responses to sotorasib-T-DXd combination on HER2 expression, supported by our preclinical studies, we evaluated HER2 expression in a patient cohort with NSCLC. The patient cohort in our study predominantly consisted of older individuals with advanced-stage adenocarcinoma, a majority of whom had negative PD-L1 expression and *KRAS* mutations. This demographic aligns with the typical clinical profile of patients with advanced NSCLC, particularly those eligible for targeted therapies. The high prevalence of driver mutations, especially *KRAS*, underscores the importance of exploring targeted treatment options in this patient population. While multiple studies have assessed HER2 expression by IHC in NSCLC^,31,32^ our study is unique in directly comparing two interpretation guidelines (developed for BC and GEA) within this context. Additionally, it is distinguished by the inclusion of multiple clinical tumor samples (pre-treatment and on-treatment) as well as post-treatment primary and metastatic tumor samples from each patient, collected through the Moffitt Rapid Tissue Donation Program.

In our cohort, 8% and 10% of the samples showed IHC score of 3+ according to BC and GEA guidelines, respectively. In a phase II clinical trial evaluating the efficacy and safety of T-DXd in patients with HER2-expressing solid tumors, the greatest clinical benefit was observed in the HER2 3+ group.^33^ This clinical trial was one of three pivotal studies that led to the Food and Drug Administration’s approval of fam-trastuzumab deruxtecan-nxki on April 5, 2024, for adult patients with unresectable or metastatic HER2-positive (IHC 3+) solid tumors who have previously received systemic treatment and have no satisfactory alternative options.^34^ In the clinical trials leading to that approval, the interpretation guideline developed for GEA was used to assess HER2 expression. It is important to consider the possibility that tumors classified as IHC 3+ by the GEA guideline could be classified as IHC 2+ by the BC guideline. This discrepancy arises due to assessing only complete membranous positivity according to BC interpretation guideline, while GEA interpretation guideline recommends to assess both complete and incomplete membranous positivity.^25,27^ In our study the discrepancy between the two interpretation guidelines was observed in 13% of the patient samples. This discrepancy in between interpretation guidelines underscores the challenges of accurately assessing HER2 expression in NSCLC using existing interpretation guidelines and highlighting the need for a new interpretation guideline for NSCLC.

In our cohort, 12% and 20% of tumors were scored as IHC 2+ using the BC and GEA interpretation guidelines, respectively, while 10% and 2% of tumors were scored as IHC 1+ using the BC and GEA interpretation guidelines, respectively. It is well established that patients with BC or GEA showing high HER2 expression (3+ or 2+/ISH amplified) experience the greatest survival benefit from HER2-targeted therapies. However, recent studies showed that patients with HER2-low phenotype (2+/ISH non-amplified or 1+) BC may also benefit from T-DXd.^17^ Studies have reported that, in addition to patients with HER2-low BC, those with other solid tumors exhibiting a HER2-low phenotype may also derive clinical benefit from T-DXd.^33^ The incidence of lung tumors showing HER2-low phenotype was reported as 46.9% in previous studies.^35^The presence of clinical benefit of T-DXd in HER2-low tumors indicates the potential utility of including HER2-low NSCLC in future studies exploring the clinical benefit of T-DXd alone or in combination with other therapies.^33,35,36^

Regarding HER2 IHC interpretation, only membranous HER2 expression has been evaluated for therapeutic decision-making in clinical care. Interestingly, nuclear HER2 expression was detected in 25 samples (13%) in this study, including two samples from two patients with *KRAS*-mutant NSCLC where an increased membranous HER2 expression (IHC score of 2+) accompanied a positive HER2 nuclear expression. Although nuclear HER2 expression has been identified as an independent prognostic factor or a contributing factor to drug resistance mechanisms in BC,^37,38^ the clinical significance of nuclear HER2 expression in NSCLC requires further investigation.

Our analysis comparing HER2 expression in KRAS-mutant NSCLC to NSCLC with other driver genomic alterations revealed a trend toward higher HER2 expression in KRAS-mutant tumor samples. However, this difference did not reach statistical significance, potentially due to the limited sample size or the absence of a true biological difference. Larger studies are warranted to clarify whether HER2 expression is indeed elevated in KRAS-mutant NSCLC compared to other genomic subtypes.

Our results also highlight the heterogeneity of HER2 expression in NSCLC and the potential for HER2-directed therapies even in patients without HER2 mutations. We observed significant differences in HER2 expression between clinical and autopsy samples. Clinical samples exhibited a higher prevalence of HER2 expression, suggesting that HER2 expression may diminish over time, during treatment, or in the context of disease progression. Although adequate preservation of protein antigenicity in rapid autopsy samples has been shown previously,^21^ it is important to consider the potential effects of decreased antigenicity and reduced IHC positivity in postmortem rapid autopsy samples compared to clinical samples. These differences may not reflect true biological variation but could instead arise from autolysis in autopsy samples, a process that inherently begins after death.^39^

A significant aspect of our study was the comparison of two HER2 IHC antibody clones, 4B5 (FDA approved-IVD) versus EP1045Y (RUO). Our results indicate that the RUO antibody detected a substantially higher percentage of HER2-positivity than the IVD one, with significant discrepancies observed in both BC and GEA interpretation guidelines. Given these findings, it is crucial to validate and standardize interpretation of HER2 protein expression in NSCLC, as the choice of IHC antibody clone could significantly influence treatment decisions and outcomes.

Due to discrepancies noted in HER2 IHC results between BC and GEA interpretation guidelines, we conducted ISH analysis in 38 samples. The ISH analysis revealed that *HER2* gene amplification or polysomy does not always accompany increased HER2 protein expression detected by IHC, consistent with previous findings in NSCLC.^40^ The observation of high HER2 expression in the absence of gene amplification points to non-genetic mechanisms such as transcriptional or post-transcriptional regulation. Indeed, it has been demonstrated that BCs showing high HER2 protein expression by IHC, but lacking gene amplification by ISH, can still respond to anti-HER2 therapies.^41^ However, the nature of the mechanism of high HER2 protein expression might influence the long-term responses, as gene amplification might restrict the ability of neoplastic populations to adapt to therapeutic pressures. While incorporating ISH into HER2 interpretation may provide additional clarity in cases with equivocal (2+) or discordant IHC results, the efficacy of anti-HER2 treatments in NSCLC patients with high HER2 protein expression without corresponding gene amplification remains an area requiring further investigation. Thus, our results highlight the complexity of HER2 regulation in NSCLC and emphasize the importance of comprehensive testing to fully capture the spectrum of HER2 alterations.

In conclusion, our study highlights the complexity of HER2 expression in NSCLC and the potential for combining HER2-directed therapies with *KRAS*^G12C^i to improve outcomes in *KRAS^G12C^*-mutant tumors. The variability in HER2 IHC scores across different interpretation guidelines, different antibody clones and clinical vs. autopsy samples underscores the need for standardization in HER2 testing. These insights lay the groundwork for future studies and underscore the need for comprehensive HER2 testing to fully capture the therapeutic potential of HER2-directed treatments in NSCLC.

## Acknowledgments

We would like to thank Angela Todica for helping us procure TDX-d, Amer A. Beg for securing grants and funding, and Alberto A. Chiappori, Andreas N. Saltos, Benjamin C. Creelan, Charles C. Williams, George Simon, Jhanelle E. Gray, Michael R. Shafique, Scott J. Antonia, and Tawee Tanvetyanon for their efforts in recruiting patients, as well as Rob Macaulay for helping autopsies for the RTD program. We would like to extend our heartfelt appreciation to the patients and their loved ones who have graciously provided consent for participation in the RTD.

## Study approval

This work has been carried out in accordance with The Code of Ethics of the World Medical Association (Declaration of Helsinki) for experiments involving humans. Tissue samples were procured in line with WHO Guiding Principles on Human Cell, Tissue and Organ Transplantation. Moffitt Cancer Center (MCC) Thoracic Rapid Tissue Donation (RTD) program was approved by the institutional review board (IRB, Advarra, Columbia, MD, Pro00030829).

## Research Data/Data Availability

The data that support the findings of this study are available from the corresponding authors upon reasonable request.

## Author contributions

H.O., B.D., A.M., B.P. and E.B.H. conceptualized the project; E.B.H, and T.A.B obtained funding; G.S.N. consented the patients; H.O., T.A.B., and H.T.B. performed autopsies; B.D., M.H., H.S., and A.M. conducted pre-clinical studies; H.O. and H.B. reviewed IHC slides; D.K. and D.C. analyzed data and performed statistical analyses; H.O., G.S.N. and B.P. characterized the clinical data. H.O., B.D., A.M., and B.P drafted the manuscript. All authors read and approved the final manuscript.

## Declaration of Generative AI and AI-assisted technologies in the writing process

During the preparation of this work the author(s) used ChatGPT to improve readability and language. After using this tool/service, the author(s) reviewed and edited the content as needed and take(s) full responsibility for the content of the publication.

## Funding

The Rapid Tissue Donation (RTD) protocol was funded by Bristol Myers Squibb, and both RTD protocol and this study were funded by the Moffitt Lung Cancer Center of Excellence. The Tissue Core, and Biostatistics and Bioinformatics Shared Resource are funded in part by the NCI Cancer Center Support grant, which confers Moffitt’s status as an NCI designated Comprehensive Cancer Center (NCI P30-CA076292). Preclinical studies were supported by the Moffitt Lung Cancer Center of Excellence and by NIH UO1 CA280829.

## Conflict of Interest

Hilal Ozakinci: No competing interest to disclose related to this work.

Bina Desai: No competing interest to disclose related to this work.

Denise Kalos: No competing interest to disclose related to this work.

Dung-Tsa Chen: No competing interest to disclose related to this work.

Menkara Henry: No competing interest to disclose related to this work.

Hitendra Solanki: No competing interest to disclose related to this work.

Theresa A. Boyle: No competing interest to disclose related to this work.

Humberto E. Trejo Bittar: No competing interest to disclose related to this work.

Gina S. Nazario: No competing interest to disclose related to this work.

Eric B. Haura: Receives research support from Revolution Medicines. E.B.H is doing advisory board work with Amgen, Janssen and RevMed and consulting for Ellipses, Kanaph Therapeutics and ORI Capital II.

Andriy Marusyk: No competing interest to disclose related to this work.

Bruna Pellini: Receives research support to the institution from Merck, has received research support to the institution from Bristol Myers Squibb, speaker honoraria from AstraZeneca, Merck, Foundation Medicine, Regeneron, and has done consulting/advisory board work with AstraZeneca, Bayer, Bristol Myers Squibb, Catalyst, Gilead, Guardant Health, Foundation Medicine, Illumina, Regeneron, Merus, and Oncohost. B.P. reports funding to the institution from the Bristol Myers Squibb Foundation/the Robert A. Winn Diversity in Clinical Trials Awards Program.

## Supplementary Materials

**Supplementary Figure 1.**
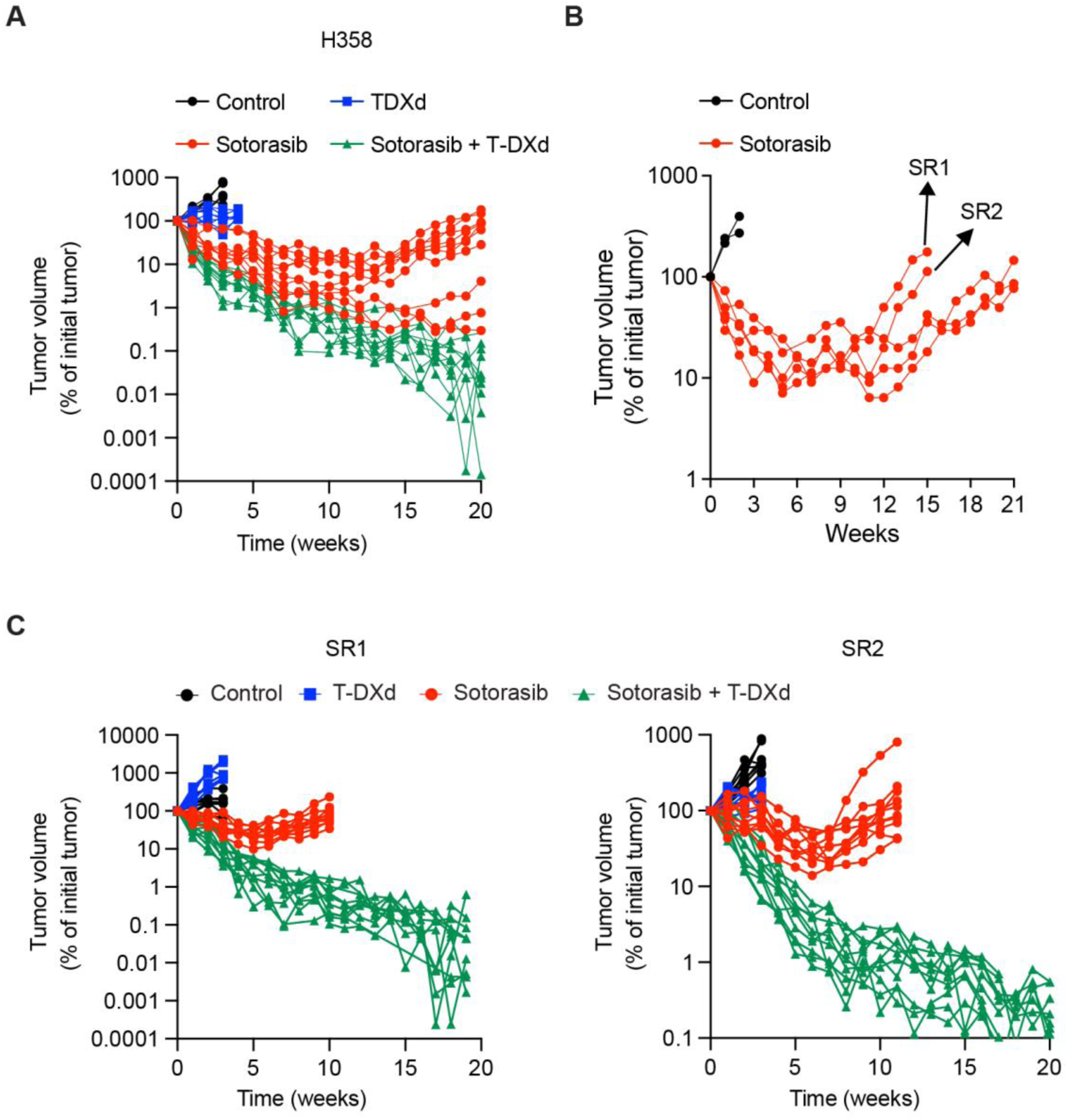
**A.** Volumetric traces of individual H358 xenograft tumors treated with vehicle control, 50 mg/kg sotorasib, 10 mg/kg T-DXd, and sotorasib/T-DXd combinations. **B.** Volumetric traces of H358 xenograft tumors treated with vehicle control, 50 mg/kg sotorasib, arrows indicate the in vivo derived sotorasib resistant models, SR1 and SR2. **C**. Volumetric traces of individual xenograft tumors of SR1 and SR2 treated with vehicle control, 50 mg/kg sotorasib, 10 mg/kg T-DXd, and sotorasib/T-DXd combinations.

**Supplementary Figure 2.**
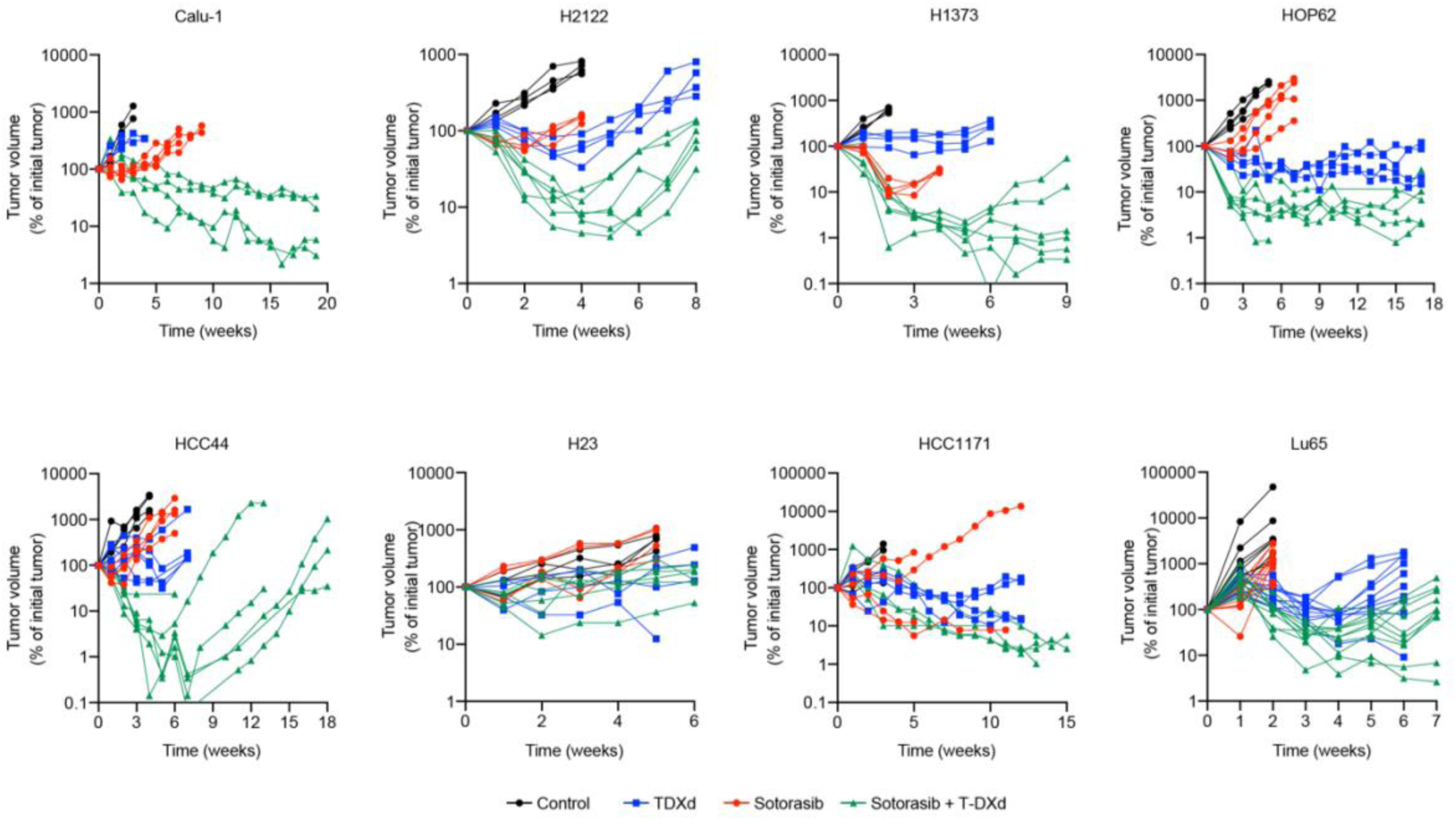
Response dynamics of individual tumors of the indicated KRAS^G12C^ cell line derived xenograft models treated with vehicle control, 50 mg/kg sotorasib, 10 mg/kg T-DXd, and sotorasib/T-DXd combinations.

**Supplemental Figure 3.**
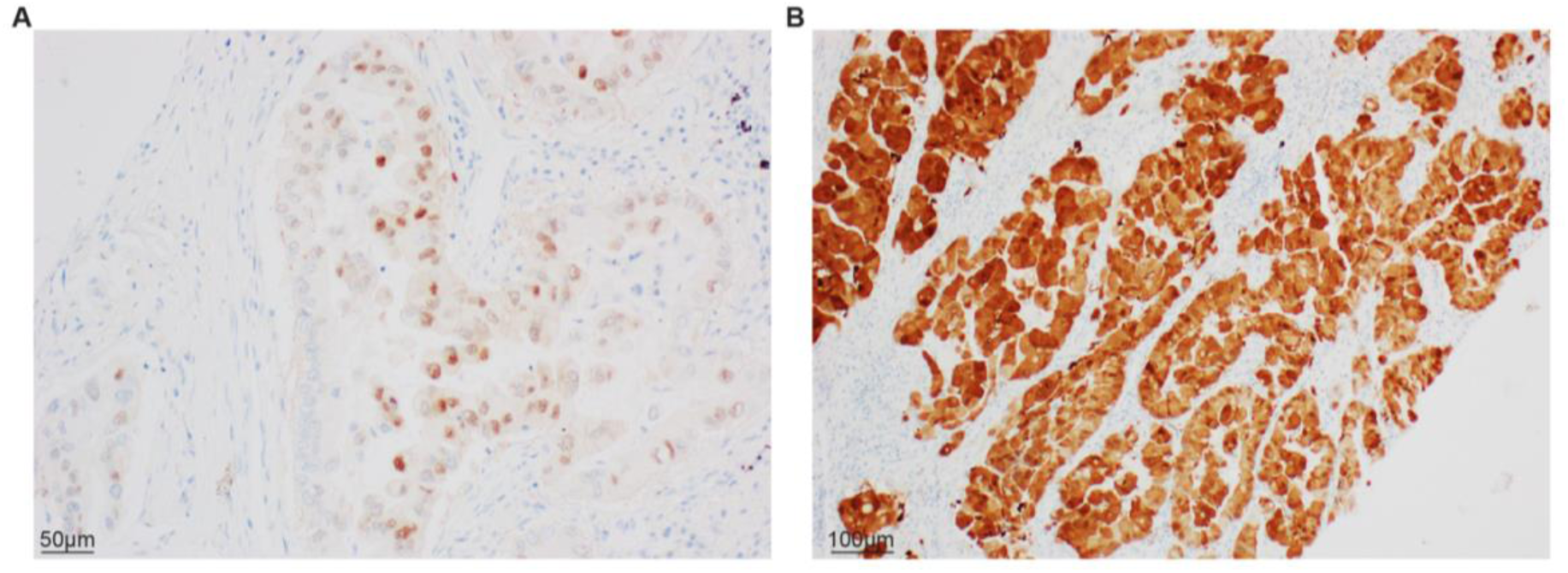
Unexpected nuclear localization of HER2 protein in the lung tumor of a patient with *KRAS^G12C^*-mutant NSCLC. **A.** IHC-stained section revealing nuclear localization of HER2 protein. **B.** IHC-stained section from different part of the tumor showing both nuclear and intense cytoplasmic HER2 staining (x10).

**Supplemental Figure 4.**
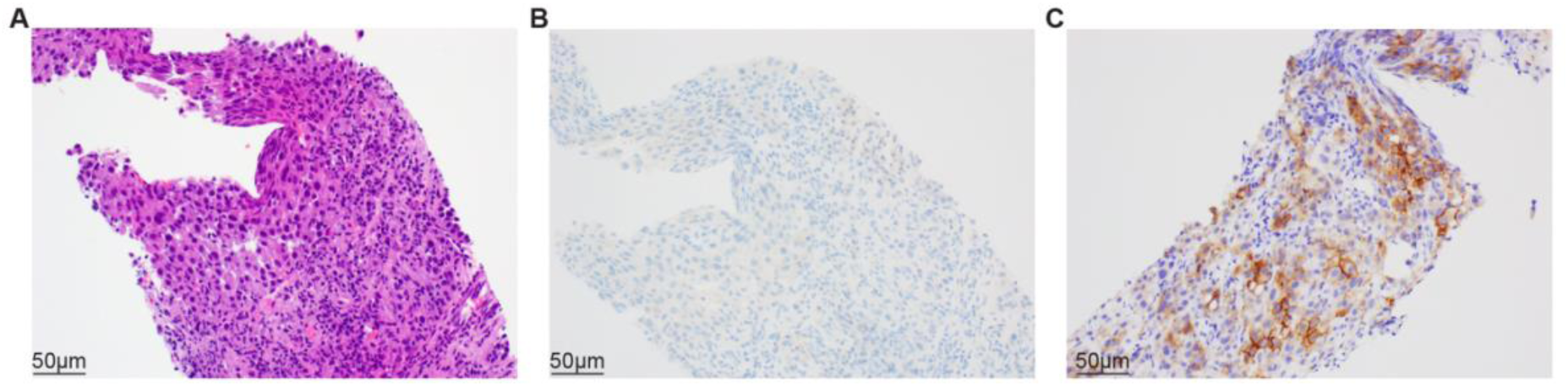
HER2 Immunohistochemistry (IHC) scoring discrepancies observed between two different antibody clones (IVD vs RUO). **A.** Representative hematoxylin and eosin (H&E) stained section depicting a poorly differentiated metastatic adenocarcinoma of the lung in the adrenal gland (x20). **B.** IHC-stained section using clone 4B5 (IVD), illustrating absence of membranous activity (score: 0) in accordance with ASCO/CAP guidelines (x20). **C**.IHC-stained section using clone EP1045Y (RUO), demonstrating complete membranous reactivity (score: 3) (x20).

**Supplemental Figure 5.**
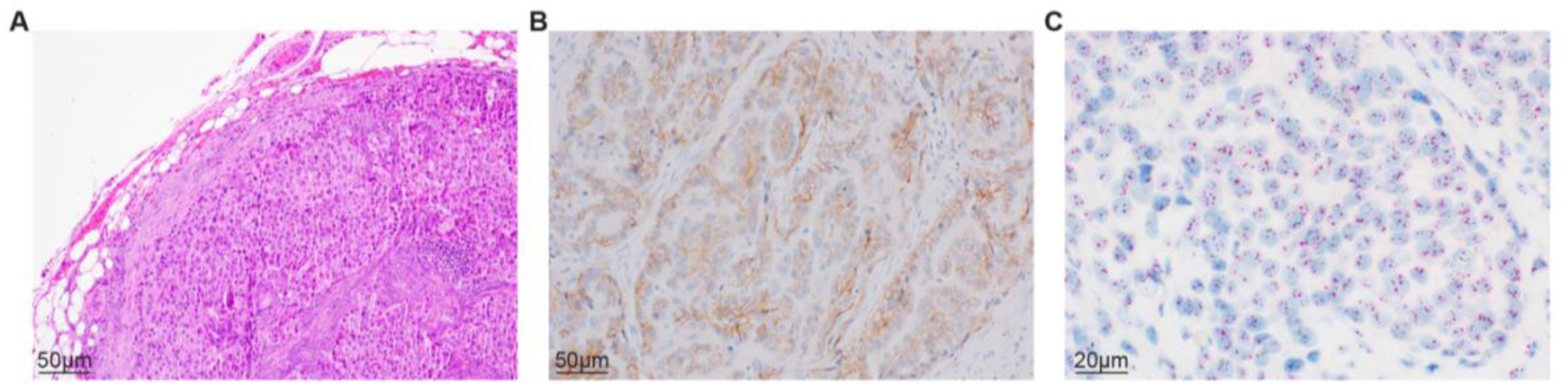
Assessment of HER2 gene amplification in a metastatic tumor with equivalent HER2 IHC score. **A.** Hematoxylin and eosin (H&E) stained section showing a poorly differentiated metastatic adenocarcinoma of the lung within a lymph node (x10). **B.** IHC-stained section displaying a 2+ (equivocal) HER2 IHC score with both breast cancer and gastroesophageal adenocarcinoma interpretation guidelines (clone: 4B5, x20). **C.** A dual in-situ hybridization (DISH) image, showing mean number of HER2 signals per cell 4.5, CEP17 signals per cell 3.7, and a HER2/CEP17 ratio of 1.21. According to ASCO/CAP combined IHC and DISH interpretation guideline, additional work-up (e.g. recounting by another pathologist) is needed to determine HER2 status (x40).

**Supplemental Table 1.** American Society of Clinical Oncology and College of American Pathologists interpretation guidelines of HER2 IHC for breast cancer (March 2023), gastroesophageal adenocarcinoma (June 2017)

**Supplemental Table 2.** Comparison of HER2 IHC scores in NSCLC patients with driver mutation versus NSCLC with no driver mutation.

**Supplemental Table 3.** Comparison of HER2 IHC scores according to BC interpretation guideline in patients with *KRAS*-mutant NSCLC versus NSCLC with other drivers.

**Supplemental Table 4.** Comparison of HER2 IHC scores in patients with *KRAS*-mutant NSCLC versus NSCLC with no driver mutation.

**Supplemental Table 5.** Comparison of HER2 IHC scores in patients with *KRAS^G12C^*-mutant NSCLC versus *KRAS^non-G12C^*-mutant NSCLC.

**Supplemental Table 6.** Comparison of HER2 IHC scores in pre-treatment (clinical) versus post-treatment (autopsy) samples from the same tumor.

**Supplemental Table 7.** Comparison of HER2 IHC scores in between IVD (4B5) and RUO (EP1045Y) clones across 26 samples.

**Supplemental Table 8.** HER2 IHC scores and DISH results for samples with discordant IHC scores or for samples with an IHC score of 2+ using both interpretation guidelines.

