## Supplemental Tables for "Trastuzumab Deruxtecan Combination Strongly Enhances Responses and Overcomes Sotorasib Resistance in *KRAS*^G12C^-Mutant NSCLC"

**Supplemental Table 1.** American Society of Clinical Oncology and College of American Pathologists interpretation guidelines of HER2 IHC for breast cancer (March 2023), gastroesophageal adenocarcinoma (June 2017)

| HER2 IHC Score | Breast cancer | Gastroesophageal adenocarcinoma | | HER2 Expression Assessment |
|---|---|---|---|---|
| | | Surgical Specimen | Biopsy Specimen | |
| 0 | No staining is observed or Membrane staining that is incomplete and is faint/barely perceptible and in ≤10% of tumor cells | No reactivity or membranous reactivity in <10% of cancer cells | No reactivity or no membranous reactivity in any cancer cell | Negative by IHC |
| 1+ | Incomplete membrane staining that is faint/ barely perceptible and in >10% of tumor cells | Faint or barely perceptible membranous reactivity in ≥10% of cancer cells; cells are reactive only in part of their membrane | Cancer cell cluster** with a faint or barely perceptible membranous reactivity irrespective of percentage of cancer cells positive | Negative by IHC |
| 2+ | Weak to moderate complete membrane staining observed in >10% of tumor cells* | Weak to moderate complete, basolateral or lateral membranous reactivity in ≥10% of tumor cells | Cancer cell cluster** with a weak to moderate complete, basolateral, or lateral membranous reactivity irrespective of percentage of cancer cells positive | Equivocal by IHC[#] |
| 3+ | Circumferential membrane staining that is complete, intense and in >10% of tumor cells | Strong complete, basolateral or lateral membranous reactivity in ≥10% of cancer cells | Cancer cell cluster** with a strong complete basolateral, or lateral membranous reactivity irrespective of percentage of cancer cells positive | Positive |
| | *Circumferential membrane IHC staining that is intense but within ≤10% of | | ** Cancer cell cluster consisting of ≥5 neoplastic cells. | [#]Must order reflex test; same specimen using ISH or order a new test if new specimen available using IHC or ISH |

| | |
|---|---|
| | tumor cells (heterogeneous but very limited in extent) can be considered 2+. |

| **Supplemental Table 2.** Comparison of HER2 IHC scores in NSCLC patients with driver mutation versus NSCLC with no driver mutation. | | | |
|---|---|---|---|
| | According to interpretation guideline for BC | | |
| | Any driver*, N=27 | No driver, N=4 | p-value |
| Patients with at least one sample showing 2+ and/or 3+ HER2 IHC score, No. (%) | 12 (44%) | 1 (25%) | >0.9 |
| | According to interpretation guideline for GEA | | |
| | Any driver*, N=27 | No driver, N=4 | p-value |
| Patients with at least one sample showing 2+ and/or 3+ HER2 IHC score, No. (%) | 15 (56%) | 2 (50%) | 0.8 |
| | ******ALK,EGFR, KRAS, MET, NRAS,ROS1* | | |

| **Supplemental Table 3**. Comparison of HER2 IHC scores according to BC interpretation guideline in patients with KRAS-mutant NSCLC versus NSCLC with other drivers. | | | |
|---|---|---|---|
| | *KRAS*, N=17 | Other drivers*, N=10 | p-value |
| Patients with at least one sample showing 2+ and/or 3+ HER2 IHC score, No. (%) | 10 (59%) | 2 (20%) | >0.9 |
| | | **ALK,EGFR, MET, NRAS,ROS1* | |

| **Supplemental Table 4**. Comparison of HER2 IHC scores in patients with KRAS-mutant NSCLC versus NSCLC with no driver mutation. | | | |
|---|---|---|---|
| | According to interpretation guideline for BC | | |
| | *KRAS*, N=17 | None, N=4 | p-value |
| Patients with at least one sample showing 2+ and/or 3+ HER2 IHC score, No. (%) | 10 (59%) | 1 (25%) | >0.9 |
| | According to interpretation guideline for GEA | | |
| | *KRAS*, N=17 | None, N=4 | p-value |
| Patients with at least one sample showing 2+ and/or 3+ HER2 IHC score, No. (%) | 11 (65%) | 2 (50%) | 0.9 |

| **Supplemental Table 5**. Comparison of HER2 IHC scores in patients with KRASG12C-mutant NSCLC versus KRASnon-G12C-mutant NSCLC. | | | |
|---|---|---|---|
| | According to interpretation guideline for BC | | |
| | *KRAS G12C*, N=9 | *KRAS* Non-G12C, N=8 | p-value |
| Patients with at least one sample showing 2+ and/or 3+ HER2 IHC score, No. (%) | 5 (56%) | 5 (63%) | 0.6 |
| | According to interpretation guideline for GEA | | |
| | *KRAS G12C*, N=9 | *KRAS* Non-G12C, N=8 | p-value |
| Patients with at least one sample showing 2+ and/or 3+ HER2 IHC score, No. (%) | 6 (67%) | 5 (63%) | 0.8 |

**Supplemental Table 6**. Comparison of HER2 IHC scores in pre-treatment (clinical) versus post-treatment (autopsy) samples from the same tumor.

| **Case #** | **Driver mutation** | **Targeted therapy** | **Tumor site/sites** | **HER2 IHC score change after treatment** |
|---|---|---|---|---|
| 1 | *KRAS* G12V | None | Lung | Decreased (positive to negative) |
| 9 | *EGFR* ex20 insertion | Abemaciclib (clinical trial) | Lung & lymph node | Unchanged (negative) |
| 16 | *KRAS* G12V | None | Lung | Decreased (positive to negative) |
| 34 | None | None | Lung | Unchanged (negative) |
| 36 | *KRAS* G12C | Sotorasib | Lung | Decreased (positive to negative) |
| 44 | *KRAS* G12C | Sotorasib | Lung | Unchanged (negative) |
| 45 | *KRAS* G12C | Sotorasib | Soft tissue | Increased (negative to equivocal) |
| 54 | *ROS1* fusion | Repotrectinib, crizotinib | Lymph node | Increased (equivocal to positive) |
| 55 | None | None | Lung&lung | Unchanged (negative) |
| *HER2 IHC scores for breast cancer and gastroesophageal adenocarcinoma algorithms were consistent across all samples. | | | | |

| **Supplemental Table 7**. Comparison of HER2 IHC scores in between IVD (4B5) and RUO (EP1045Y) clones across 26 samples. | | | | | |
|---|---|---|---|---|---|
| According to interpretation guideline for BC | | | | | |
| | | HER2 IHC score with IVD clone | | | |
| | | 3+ | 2+ | 0 or 1+ | Total |
| HER2 IHC score with research clone | 3+ | 0 (0%) | 1 (3.8%) | 1 (3.8%) | 2 (7.7%) |
| | 2+ | 0 (0%) | 1 (3.8%) | 12 (46%) | 13 (50%) |
| | 0 or 1+ | 0 (0%) | 0 (0%) | 11 (42%) | 11 (42%) |
| | Total | 0 (0%) | 2 (7.7%) | 24 (92%) | 26 (100%) |
| According to interpretation guideline for GEA | | | | | |
| | | HER2 IHC score with IVD clone | | | |
| | | 3+ | 2+ | 0 or 1+ | Total |
| HER2 IHC score with research clone | 3+ | 0 (0%) | 1 (3.8%) | 1 (3.8%) | 2 (7.7%) |
| | 2+ | 0 (0%) | 2 (7.7%) | 14 (54%) | 16 (62%) |
| | 0 or 1+ | 0 (0%) | 0 (0%) | 8 (31%) | 8 (31%) |
| | Total | 0 (0%) | 3 (12%) | 23 (88%) | 26 (100%) |

| **Supplemental Table 8**. HER2 IHC scores and DISH results for samples with discordant IHC scores or for samples with an IHC score of 2+ using both interpretation guidelines. | | | | | | | | | | |
|---|---|---|---|---|---|---|---|---|---|---|
| Case # | Driver mutation | Sample type | Tumor site | IHC scores | | DISH assessment | | | Combined IHC and DISH Interpretation | |
| | | | | BC | GEA | Average *HER2* gene copy number | Average CEP17 copy number | *HER2/CEP17* ratio | Breast cancer | Gastroesophageal adenocarcinoma |
| 3 | KRAS | Autopsy | Lung | 2 | 3 | 5.25 | 2.25 | 2.3 | IHC: equivocal, ISH: positive | IHC:positive, ISH:positive |
| 13 | ALK | Autopsy | Pleura | 1 | 2 | 4.15 | 2.1 | 1.97 | IHC: negative, ISH: additional work-up | IHC:equivocal, ISH:negative |
| 16 | KRAS | Clinical | Liver | 2 | 3 | 5.6 | 3.1 | 1.8 | IHC: equivocal, ISH: additional work-up | IHC:positive, ISH:additional work-up |

| | | | | | | | | | | |
|---|---|---|---|---|---|---|---|---|---|---|
| 16 | KRAS | Autopsy | Lymph Node | 2 | 3 | 5.7 | 0.9 | 1.16 | IHC: equivocal, ISH: additional work-up | IHC:positive, ISH:negative |
| 16 | KRAS | Autopsy | Lymph Node | 1 | 2 | 6.35 | 5.55 | 1.14 | IHC: negative, ISH: additional work-up | IHC:equivocal, ISH:positive |
| 16 | KRAS | Autopsy | Lymph Node | 1 | 2 | 4.6 | 4.4 | 1.04 | IHC: negative, ISH: additional work-up | IHC:equivocal, ISH:additional work-up |
| 16 | KRAS | Autopsy | Soft tissue | 1 | 2 | 4.2 | 4.8 | 0.87 | IHC: negative, ISH: additional work-up | IHC:equivocal, ISH:additional work-up |
| 21 | KRAS | Autopsy | Brain | 2 | 3 | 5.05 | 2.85 | 1.77 | IHC: equivocal, ISH: additional work-up | IHC:positive, ISH:negative |
| 28 | MET | Autopsy | Lung | 1 | 2 | 2.3 | 1.85 | 1.24 | IHC: negative, ISH: negative | IHC:equivocal, ISH:negative |
| 28 | MET | Autopsy | Lung | 1 | 2 | 2.95 | 2.15 | 1.37 | IHC: negative, ISH: negative | IHC:equivocal, ISH:negative |
| 28 | MET | Autopsy | Lymph node | 1 | 2 | 2.45 | 1.9 | 1.28 | IHC: negative, ISH: negative | IHC:equivocal, ISH:negative |
| 28 | MET | Autopsy | Bone | 1 | 2 | 3.8 | 2.25 | 1.68 | IHC: negative, ISH: negative | IHC:equivocal, ISH:negative |
| 30 | None | Autopsy | Lung | 1 | 2 | 3.3 | 2.3 | 1.43 | IHC: negative | IHC:equivocal, ISH:negative |

| | | | | | | | | | | |
|---|---|---|---|---|---|---|---|---|---|---|
| | | | | | | | | | , ISH: negative | |
| 30 | None | Autopsy | Lung | 1 | 2 | 2.45 | 1.9 | 1.28 | IHC: negative, ISH: negative | IHC:equivocal, ISH:negative |
| 30 | None | Autopsy | Lung | 1 | 2 | 2.8 | 1.85 | 1.51 | IHC: negative, ISH: negative | IHC:equivocal, ISH:negative |
| 35 | KRAS | Autopsy | Liver | 2 | 2 | 4 | 3.05 | 1.31 | IHC: equivocal, ISH: additional work-up | IHC:equivocal, ISH:additional work-up |
| 36 | KRAS | Clinical | Lymph node | 2 | 2 | 2.7 | 1.85 | 1.45 | IHC: equivocal, ISH: negative | IHC:equivocal, ISH:negative |
| 37 | KRAS | Autopsy | Lung | 2 | 2 | 3.45 | 2.7 | 1.27 | IHC: equivocal, ISH: negative | IHC:equivocal, ISH:negative |
| 37 | KRAS | Autopsy | Lung | 1 | 2 | 2.75 | 2.25 | 1.22 | IHC: negative, ISH: negative | IHC:equivocal, ISH:negative |
| 40 | KRAS | Clinical | Lung | 1 | 2 | 2.45 | 1.65 | 1.48 | IHC: negative, ISH: negative | IHC:equivocal, ISH:negative |
| 43 | EGFR | Autopsy | Lung | 2 | 2 | 2.9 | 1.75 | 1.65 | IHC: equivocal, ISH: negative | IHC:equivocal, ISH:negative |
| 43 | EGFR | Autopsy | Lymph Node | 1 | 2 | 3.05 | 2.3 | 1.32 | IHC: negative, ISH: negative | IHC:equivocal, ISH:negative |
| 43 | EGFR | Autopsy | Lymph Node | 1 | 2 | 3.65 | 2.85 | 1.28 | IHC: negative, ISH: negative | IHC:equivocal, ISH:negative |
| 43 | EGFR | Autopsy | Soft Tissue | 2 | 2 | 4 | 2.1 | 1.9 | IHC: equivocal, ISH: addition | IHC:equivocal, ISH:negative |

| | | | | | | | | | | |
|---|---|---|---|---|---|---|---|---|---|---|
| | | | | | | | | | al work-up | |
| 43 | EGFR | Autopsy | Lung | 2 | 2 | 2.85 | 2 | 1.42 | IHC: equivocal, ISH: negative | IHC:equivocal, ISH:negative |
| 43 | EGFR | Autopsy | Lung | 2 | 2 | 3.25 | 1.85 | 1.75 | IHC: equivocal, ISH: negative | IHC:equivocal, ISH:negative |
| 45 | KRAS | Autopsy | Bone/Soft tissue | 2 | 2 | 2.55 | 2 | 1.2 | IHC: equivocal, ISH: negative | IHC:equivocal, ISH:negative |
| 45 | KRAS | Autopsy | Lymph Node | 2 | 2 | 2.5 | 1.9 | 1.31 | IHC: equivocal, ISH: negative | IHC:equivocal, ISH:negative |
| 45 | KRAS | Autopsy | Lymph Node | 2 | 2 | 2.6 | 1.75 | 1.48 | IHC: equivocal, ISH: negative | IHC:equivocal, ISH:negative |
| 52 | None | Autopsy | Adrenal | 2 | 2 | 4.55 | 3.3 | 1.37 | IHC: equivocal, ISH: additional work-up | IHC:equivocal, ISH:additional work-up |
| 52 | None | Autopsy | Liver | 1 | 2 | 3.85 | 3.05 | 1.26 | IHC: negative, ISH: negative | IHC:equivocal, ISH:negative |
| 52 | None | Clinical | Adrenal | 1 | 2 | 5.15 | 3.85 | 1.33 | IHC: negative, ISH: additional work-up | IHC:equivocal, ISH:additional work-up |
| 52 | None | Autopsy | Liver | 1 | 2 | 3.85 | 3.3 | 1.16 | IHC: negative, ISH: negative | IHC:equivocal, ISH:negative |
| 52 | None | Autopsy | Liver | 1 | 2 | 3.25 | 2.8 | 1.16 | IHC: negative, ISH: negative | IHC:equivocal, ISH:negative |
| 52 | None | Autopsy | Adrenal | 2 | 2 | 3.85 | 3.2 | 1.2 | IHC: equivoc | IHC:equivocal, ISH:negative |

| | | | | | | | | | | |
|---|---|---|---|---|---|---|---|---|---|---|
| | | | | | | | | | al, ISH: negative | |
| 52 | None | Clinical | Adrenal | 0 | 2 | 4.5 | 3.15 | 1.42 | IHC: negative, ISH: additional work-up | IHC:equivocal, ISH:additional work-up |
| 54 | ROS1 | Clinical | Pleural fluid | 2 | 3 | 3.9 | 2.9 | 1.34 | IHC: equivocal, ISH: negative | IHC:positive, ISH:negative |
| 54 | ROS1 | Clinical | Lymph Node | 2 | 2 | 4.5 | 3.7 | 1.21 | IHC: equivocal, ISH: additional work-up | IHC:equivocal, ISH:additional work-up |
